# Exploration of the Metabolic Potential of the *Corallococcus* Genus: A Rich Source of Secondary Metabolites, and CAZymes

**DOI:** 10.1101/2024.10.08.616840

**Authors:** Md. Saddam Hossain, Shanjana Rahman Tuli, Nigar Fatima, Md. Tauhidul Islam Tanim, Abu Hashem

## Abstract

Different secondary metabolites take on important roles. They can also transport metals, function as sexual hormones, facilitate harmonious coexistence of microbes and living beings, and induce differentiation, besides being used as competitive weapons against other bacteria, fungi, amoebae, plants, insects, and large animals. Progress in next-generation sequencing methods, together with microbial genome sequencing, has unveiled a wealth of natural products (NPs) with yet-untapped potential. Genome mining of *Corallococcus* species and *C. exiguus* subspecies that can make secondary metabolites that are clinically important has not been studied very much. The goal of this study is to look into Corallococcus genomes’ biosynthetic gene clusters (BGCs) and carbohydrate-active enzyme gene clusters. We fully characterize BGCs from *Corallococcus* reference genome sequences that are publicly available using bioinformatic tools in addition to phylogenetic and genomic comparisons. Our results show that there is a huge range of BGCs in different *Corallococcus* genomes, but the species or subspecies level affects the ability to make NPs. Additionally, we investigated unknown and less comparable BGCs at the species level and found that *C. llansteffanensis* has more potential as a reservoir of new secondary metabolites. These insights will pave the way for the algorithmic identification of species- and subspecies-specific pathways for NP development.

## 1. Introduction

The ongoing search for new natural resources is necessary to address the global health crisis posed by antibiotic resistance. Microbial natural products (NPs) have thermal stability as well as ease of cultivation, amenability to genetic manipulation, and year-round availability that make them an attractive alternative to plant and animal sources. Because of these fascinating qualities, scientists are now more interested in investigating the enormous potential of bioproducts originating from microbiological sources [1].

Microorganisms have an ability to make a wide variety of secondary metabolites or NPs, including non-ribosomal peptides (NRPs), polyketides (PKs), ribosomally synthesized and post-translationally modified peptides (RiPPs), saccharides, alkaloids, and terpenoids. These NPs have immense potential for applications in the pharmaceutical and agricultural industries, with over 50% of Food and Drug Administration (FDA) approved drugs and 65% of current clinical drugs inspired by NPs [2]. The biosynthesis of microbial NPs is regulated by a unique set of genes grouped into clusters called biosynthetic gene clusters (BGCs).The microbial chromosomes contain these BGCs, which aid in the co-expression of the transporters, regulators, and enzymes involved in the production and secretion of NP [3]. Despite the vast potential of NPs, only a fraction has been explored. Therefore, it’s critical to locate and investigate BGCs in microbial genomes that may have hidden metabolic potential [4].

Recent developments in data mining tools, along with strong analytical and genetic instruments, have made it possible to identify NPs that were not previously identified [5]. Gaining access to genome sequence data offers important new perspectives on the distribution, diversity, and evolutionary processes of BGCs. Furthermore, the striking structural diversity seen in NPs is probably greatly influenced by the quick evolution of BGCs in comparison to other genetic components [6]. The primary method for discovering biologically and therapeutically significant natural products is the extraction of culture filtrates, but this approach falls short because a large number of chemicals remain unexplored and the overall process is time-intensive [7].

*Corallococcus*, a genus within the Deltaproteobacteria class of the Myxococcaceae family, has garnered significant attention due to the discovery of bioactive compounds. These predatory bacteria are renowned for their distinctive rippling swarm movement, coral-shaped fruiting bodies, and broad prey range [8]. Different types of *C. coralloides* have produced some interesting chemicals, such as corallopyronins A, B, and C, corallorazine A, and coralmycins A and B which are all antibacterial [9]. Notably, the ability to degrade polysaccharides into usable tri- and oligosaccharides makes them particularly attractive for the development of novel drugs and biocontrol agents using renewable sources [10].

This study aims to delve deeper into the potential of *Corallococcus* spp. and *C. exiguus*, specifically focusing on their ability to produce novel secondary metabolites. By leveraging bioinformatics tools for BGC analysis, we aim to unlock the hidden potential of these fascinating bacteria and contribute to the development of novel bioactive compounds with promising applications in various fields.

## 2. Materials and Method

### 2.1. Genomic Data Acquisition

We downloaded all the available assembly data from refseq for *C. exiguus* and one representative genome at species level from NCBI (https://www.ncbi.nlm.nih.gov/assembly) including assembly statistic report information (accessed on 21 October, 2023). We removed contigs less than 1000 bp and used ContEst16S for to check for contamination [11]. The List of Prokaryotic names with Standing in Nomenclature (LPSN) database was used to identify the number of species for *Coralococcus* (https://lpsn.dsmz.de/). Genomic data were selected if the average gene and protein number fell within a range of μ – 2σ and μ + 2σ, where μ is the average number of species protein or gene sequences and σ is the standard deviations.

### 2.2. Comparative and Evolutionary Study

The genetic diversity of genomes was assessed using pairwise average nucleotide identity (ANI) analysis, employing a Python package known as pyANI, to identify distinct species boundaries. The evolutionary relationships among all *Corallococcus* at species and subspecies level were investigated through a phylogenomic analysis based on Genome BLAST Distance Phylogeny (GBDP), utilizing the TYGS (Type Strain Genome Server) [12]. The resulting phylogenetic tree was visualized and presented using the interactive tree of life (iToL version 6.7.5). To create a comprehensive overview of each genome’s structure, a circular map was generated for each genome using the BLAST Ring Image Generator (BRIG) [13]. After that, all the protein sequences that had been labeled were analyzed using the Python-based cogclassifier 1.0.5 tool to sort them into groups based on their functional roles. To identify acquired antibiotic resistance genes and characterize pathogenicity, the Center for Epidemiology Services (https://www.genomicepidemiology.org/) resources, ResFinder and Pathogenicity, were employed.

### 2.3. Biosynthetic Gene Clusters (BGCs) Predictions

The antibiotics and secondary metabolite analysis shell (antiSMASH) online version 7.0 was used to identify potential biosynthetic gene cluster (BGC) regions, and the resulting job ID was submitted to the biosynthetic gene cluster family (BiG-FAM) database to obtain the corresponding Gene Cluster Family (GCF) classification [[14, 15]]. The antiSMASH analysis was conducted using fasta DNA as input, with a relaxed detection strictness setting. KnownClusterBlast, ClusterBlast, SubClusterBlast, and MIBiG cluster comparison were enabled as additional features.

### 2.4. BAGEL Analysis

The identification of potential bacteriocins and RiPPs was carried out using BAGEL4 [16]. The predicted antimicrobial peptides were then subjected to further analysis using iAMPRED to evaluate their antimicrobial potential and AllerTOP v. 2.0 to assess their allergenicity [17, 18].

### 2.5. Exploration of unknown and less similar BGCs

We investigated T1PKS-NRPS hybrid, NRPS, and T1PKS BGCs in *Corallococcus* species that were entirely unidentified and shared less than 50% similarity with known clusters. The sequence-based similarity network (SSN) produced by BiG-SCAPE was visualized using the Cytoscape 3.10 software [19]. We utlized Antibiotic Resistant Target Seeker (ARTS) version 2.0 and Resistance Gene Identifier (RGI) to examine BGCs for resistance genes, and subsequently used Clinker version 0.0.28 to compare and visualise the prioritised BGCs [20–22]. NaPDoS pipeline was used to compare C domain sequences with a database of characterized natural products and construct a phylogenetic tree which was visualized using iToL [23, 24]. Using PRISM version 4.4.5, we predicted the structure of compounds produced by the prioritized biosynthetic gene clusters (BGCs) [25]. Subsequently, deepFRI utilizing graph convolutional networks, was employed to predict the functions of these compounds [26]. Finally, Passonline was used to predict antimicrobial activities associated with the structures predicted by PRISM [27].

### 2.6. Exploration of Carbohydrate Active Enzymes (CAZymes)

The identification of genes encoding carbohydrate-active enzymes (CAZymes) was performed using the dbCAN3 web server, which employs DIAMOND, HMMER, CAZy, dbCAN, and dbCAN-sub databases for protein annotation. Only genes identified by at least two of the three tools were retained. Additionally, dbCAN3 was used to detect CAZyme gene clusters (CGCs) and signal peptides in all the studied *Coralloccus* genomes [28].

## 3. Results

### 3.1. Studied Genome’s General Features

The LPSN database, a comprehensive repository of microbial taxonomy listed 13 distinct *Corallococcus* species, providing valuable insights into their evolutionary origins, ecological niches, and potential biotechnological applications. There was a difference in genome size between *Corallococcus* species. *Corallococcus carmarthensis* CA043D (GCF_003611695.1) had the largest genome, with 10,776,576 base pairs. On the other hand, *Corallococcus macrosporus* DSM 14697 (GCF_002305895.1) had the smallest genome, with 8,973,512 base pairs. The average genome size across *Corallococcus* species was 9,995,176.23 base pairs. In addition to genome size diversity, variations in GC content were also observed among *Corallococcus* species. The lowest GC content, 69.56%, was in *Corallococcus sicarius* CA040B (GCF_003611735.1), while the highest GC content, reaching 70.69%, was found in *Corallococcus interemptor* AB047A (GCF_003668875.1). The average GC content across *Corallococcus* species was 70.15%. ContEst16S analysis, employed to evaluate potential contamination, revealed no contamination for *Corallococcus coralloides* DSM 2259 (GCF_000255295.1). However, for other *Corallococcus* species, the results were inconclusive. The genome size of the *C. exiguus* subspecies ranged from 10304044 bp to 10538407 bp, with an average GC content of 69.59%. ContEst16S analysis revealed contamination for *C. exiguus* NCCRE002 (GCF_017302975.1), while the results for other strains were inconclusive (**Table S1**).

### 3.2. Genomic Analysis

The range of genes for *Coralococcus* species was observed from 7198 (*Corallococcus silvisoli* c25j21) to 8707 (*Corallococcus carmarthensis* CA043D). The average number of genes was 7983 with a standard deviation of 507.367085 1. A comparison of the *Corallococcus coralloides* DSM 2259 genome to other genomes using a circular map created with BRIG version 0.95 showed gaps or regions of low similarity, suggesting variations in multiple regions (**Fig. S1**). The Genome-to-Genome Distance Calculator (GGDC) approach employs Average Nucleotide Identity (ANI), digital DNA-DNA Hybridization (dDDH) values, and percent genomic G+C content variations (cut-offs of 96%, 70%, and 0.1%, respectively) to distinguish between species and subspecies of organisms. All the studied species were distinct from each other, as shown in **Tables S2** and **S3**. Additionally, the similarity among the studied genomes exhibited clustering at an average nucleotide identity (ANI), as illustrated in **Fig. 1(a)**.

**Fig. 1.**
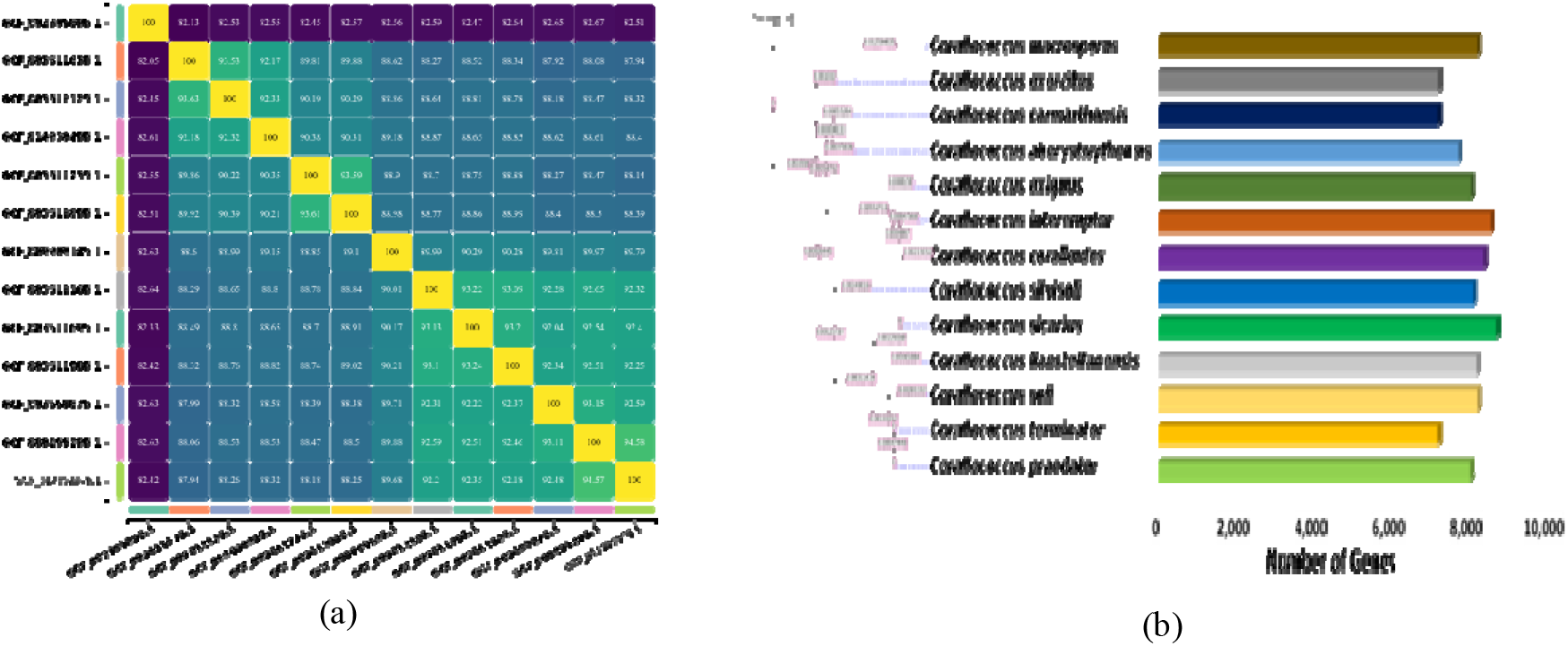
Genome comparison at *Corallococcus* species level; (a) Clustered *Corallococcus* species genomes based on their Average Nucleotide Identity (ANI) values (b) A phylogenetic tree of *Corallococcus* species, accompanied by their respective gene counts, provides valuable insights into their evolutionary relationships and gene content variations.

The range of genes across the analyzed genomes of *C. exiguus* subspecies spanned from 8,164 (GCF_009909105.1) to 8,461 (GCF_006376655.1), with an average gene count of 8,280 and a standard deviation of 589.9. Despite barely exceeding the protein count threshold, GCF_006376655.1 was retained for further analysis due to its borderline status. Subsequent CheckM and EzBiocloud analyses revealed 98.97% completeness, 2.71% contamination, and inconclusiveness respectively. Considering these findings, GCF_006376655.1 was ultimately included in our studied genomes.

The phylogenetic tree was constructed using the FastME 2.1.6.1 algorithm [29] based on Genome BLAST Distance Phylogeny (GBDP) distances derived from genome sequences. The branch lengths represent the evolutionary distances between species, as calculated using the GBDP distance formula d5 **(Fig. 1(b))**.

Twenty-five out of the 26 COG functional categories were observed in the COGs proteins of *Corallococcus* species, with the absence of the nuclear structure category **(Table S4)**. Across different *Corallococcus* species, the proportion of orthologous genes involved in secondary metabolite biosynthesis, transport, and catabolism ranged from 2.56% to 4.30%, implying a widespread presence of secondary metabolites. **(Fig. S2)**. The pangenomic analysis of *C. exiguus* using the ProPan database [30] revealed that these bacteria possess a diverse repertoire of genes involved in secondary metabolite biosynthesis, transport, and catabolism, with 176 core genes, 206 dispensable genes, and 151 unique genes **(Table S5)**. Notably, no acquired antibiotic resistance genes were detected, and all strains were considered non-pathogenic.

### 3.3. BGCs Predictions

A total of 613 BGCs were found in the 13 *Corallococcus* genomes, with an average of 47.15 BGCs per genome and a standard deviation of 12.01. The lowest number of BGCs (24) was found in *Corallococcus macrosporus* (GCF_002305895.1), while the highest number of BGCs (67) was found in *Corallococcus llansteffanensis* (GCF_003612055.1). The genome size and the number of BGCs were found to be moderately positively correlated, with a Pearson correlation coefficient of 0.6709324. The *C. exiguus* genomes, on the other hand, had a total of 601 biosynthetic gene clusters (BGCs), with an average of 50.08 BGCs per genome and a standard deviation of 8.52. The lowest number of BGCs (37) was found in GCF_017302975.1, while the highest number of BGCs (70) was found in GCF_013248925.1. The Pearson correlation coefficient between genome size and the number of BGCs was 0.16, indicating a weak positive linear relationship between genome size and BGCs. Around 22.81% of the genome at the subspecies level was made up of biosynthetic gene clusters (BGCs), which was a little higher than the 21.97% at the species level (**Table S6**).

Out of 22 biosynthetic gene clusters (BGCs) identified at the species level, eight types were present in all species. These eight types were NRPS, NRPS-T1PKS hybrid, hybrid BGCs, NRPS-like, RiPP-like, T3PKS, terpene, and thiopeptide. Among these, NRPS was the most abundant type, with a total of 155 occurrences. Similarly, out of 16 BGCs identified at the subspecies level, eight types were present in all subspecies. These eight types were NRPS, NRPS-T1PKS hybrid, hybrid, lanthipetide class II, NRPS-like, RiPP-like, T1PKS, and thiopeptide. Again, NRPS was the most abundant type, with a total of 187 occurrences (**Fig. 2(a))**.

**Fig. 2.**
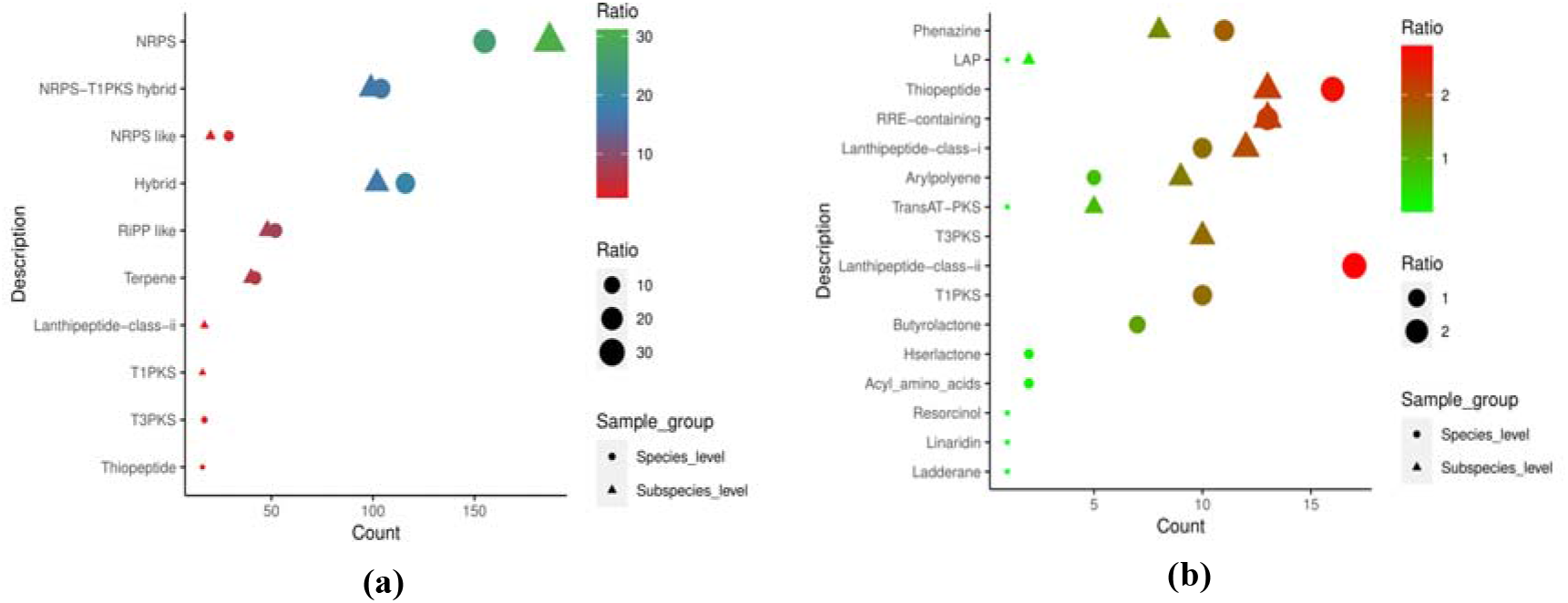
Distribution of common BGCs at both species and subspecies levels; (a) The most common BGCs (b) The less common BGCs

Acyl amino acids, Arylpolyene, Butyrolactone, Hserlactone, Ladderane, Lanthipeptide class I, Lanthipeptide class II, Linaridin, LAP, Phenazine, Resorcinol, RRE containing, T1PKS, Thiopeptide, and TransAT-PKS were less prevalent BGCs that were not present in all species. There was just one genome including BGCs for Ladderane, Linaridin, LAP, Resorcinol, and TransAT-PKS. Phenazine, LAP, Thiopeptide, RRE containing, Lanthipeptide class I, Arylpolyene, T3PKS, and TransAT-PKS were less prevalent BGCs that were absent from all subspecies. It was shown that only two subspecies had LAP BGC **(Fig. 2(b) and Fig. S4)**.

In two sets of bacteria, one at the species level and the other at the subspecies level, the prevalence and similarity of biosynthetic gene clusters (BGCs) were compared. Out of 613 BGCs identified in the species-level set, 310 (50.6%) were known, while 303 (49.4%) were unknown. Among the known BGCs, 54 (17.8%) exhibited 100% similarity, 16 (5.3%) showed similarity between 75% and 100%, 27 (8.9%) demonstrated similarity between 50% and 75%, and 213 (70.3%) exhibited less than 50% similarity. In contrast, out of 601 BGCs found in the subspecies-level set, 340 (56.6%) were known, while 261 (43.4%) were unknown. Among the known BGCs, 55 (16.2%) exhibited 100% similarity, 22 (6.5%) showed similarity between 75% and 100%, 28 (8.2%) demonstrated similarity between 50% and 75%, and 235 (69.1%) exhibited less than 50% similarity **(Table S7 and S8)**.

Out of the 613 biosynthetic gene clusters (BGCs) identified at the species level, 194 (31.65%) were classified as core BGCs, 371 (60.52%) were putative BGCs, and 48 (7.83%) were orphan BGCs based on BigFAM analysis. Core BGCs are highly conserved and are likely to be essential for the survival of the organism. Putative BGCs are less conserved and may have more recently evolved functions. Orphan BGCs are the least conserved and may have unique functions [31]. Similarly, out of the 601 BGCs identified at the subspecies level, 154 (25.62%) were classified as core BGCs, 405 (67.39%) were putative BGCs, and 42 (6.99%) were orphan BGCs based on BigFAM analysis **(Table S9)**.

### 3.4. Antimicrobial Peptide Predictions

All species except *Corallococcus praedator* (GCF_003612125.1) had at least one RiPP-encoding gene. RiPPs were split into two groups: lanthipeptides and sactipeptides. These groups have three main peptides: cerecidin, zoocin A, and fulvocin C **(Fig. S5)**. Cerecidin was further divided into three subgroups: 170.1, 171.1, and 172.1. According to iAMPRED analysis, 172.1 cerecidin, 170.1 cerecidin, and fulvocin C had antibacterial, antiviral, and antifungal activities. In contrast, 171.1 cerecidin lacks antimicrobial activity because it is not associated with a transporter protein **(Table 1)**. All of these antimicrobial peptides were predicted to be non-allergenic by the AllerTOP version 2.0 analysis. At the subspecies level, all species had three AOIs with a core peptide **(Table S10)**.

**Table 1:** Core peptides and their predicted antimicrobial capacity (values greater than 0.50 are considered prospective)

| Core Peptide Name | Antiviral | Antibacterial | Antifungal |
| --- | --- | --- | --- |
| 172.1;Cerecidin | 0.75 | 0.80 | 0.76 |
| 171.1;Cerecidin | 0.26 | 0.38 | 0.24 |
| 170.1;Cerecidin | 0.73 | 0.79 | 0.74 |
| 102.2;Fulvocin-C | 0.99 | 0.93 | 0.98 |

### 3.5. Unknown NRPS, T1PKS and T1PKS-NRPS Hybrid BGCs

There were 140 NRPS BGCs, 9 T1PKS BGCs, and 85 T1PKS-NRPS hybrid BGCs of unidentified and less similar BGCs at the Corallococcus species level. On the basis of SSN analysis, NRPS had 120 singletons and 126 families, T1PKS had 3 singletons and 4 families, and the NRPS-T1PKS hybrid had 53 singletons and 71 families (**Fig. 3**).

**Fig. 3.**
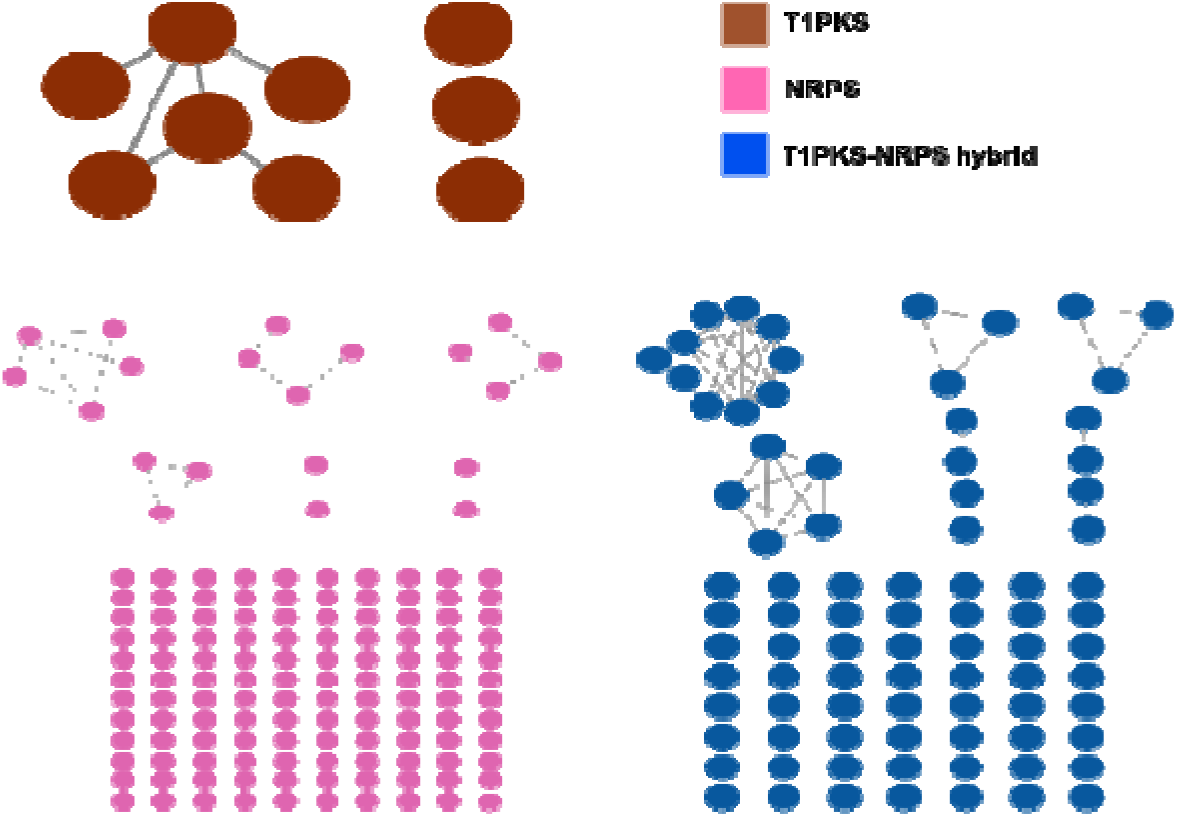
Sequence-based similarity network (SSN) of unknown and less similar NRPS, T1PKS and T1PKS-NRPS Hybrid BGCs at *Corallococcus* species level

In our study, 28 BGCs were prioritised; of these, 21 were NRPS, seven were T1PKS-NRPS hybrid, and none were T1PKS on the grounds of antibiotic resistance gene presence (**Table S11**). The mean similarity of the C domains across all prioritized NRPS BGCs was 45.21%, ranging from 28% to 53%. With a sequence similarity of greater than 90% with the biosynthetic domains of experimentally validated compounds, precise structural classification of the compounds can be anticipated from these domains [32]. This result unequivocally suggests that they may serve as a reservoir for novel biomolecules. The domains exhibited a minimal degree of sequence similarity with genes associated with biosynthetic pathways of the following compounds: Anabaenopeptilide (10), Bacitracin (2), Bleomycin (1), Fengycin (2), Lichenysin (1), Microcystin (9), Nostopeptolide (36), Pyoverdine (8), Pyridomycin (2), Surfactin (1), Tubulysin (1), and Tyrocidine (3) (**Table S12**). The product forming pathways were found in all domains but in phylogenetic analysis clustered them differently based on the sequence similarity (**Fig. 4**)

**Fig. 4.**
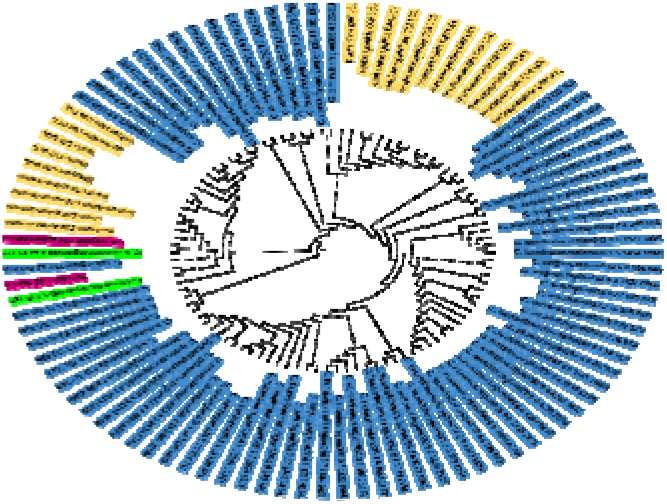
Neighbor-joining phylogenetic analysis of condensation domains of prioritized NRPS BGCs in comparison to domains in the NaPDoS database. The blue denotes the examined C domains that were unable to form clusters with any pathway. The C domains clustered with pathways (illustrated in purple) in the NaPDoS database are denoted by the color green. The yellow color represents the remaining pathways that were found in all domains.

Analysis of the clinker demonstrated that the prioritized BGCs were distinct from one another (**Fig. S6 & S7**). After conducting DeepFRI analysis, it was observed that the prioritised clusters exhibited a wide range of predicted molecular functions (MFs) and biological processes (BPs), which are indicative of their ability to produce secondary metabolites [33]. In particular, terms like “organic substance biosynthetic process,” “cellular metabolic process,” and “biosynthetic process” serve as robust indicators of the active synthesis of organic compounds (**Table S13**). With the exception of region 048, which was attributed to *Corallococcus praedator*, PRISM successfully predicted structures for every region of the prioritised NRPS BGCs **(Fig S8)**. According to the findings of Passonline analysis, the principal activities exhibited by the prioritised BGCs were antibacterial, in addition to antifungal and antiviral properties (**Table S14**)

### 3.6. CAZymes Prediction

In the Corallococcus species we examined, 2886 genes were responsible for the synthesis of carbohydrate-active enzymes (CAZYmes). These CAZYmes were classified into six families: auxiliary activities (AA), carbohydrate-binding modules (CBM), carbohydrate esterases (CE), glycoside hydrolases (GH), glycosyl transferases (GT), and polysaccharide lyases (PL) **(Table S15)**. The distribution of CAZymes across the studied *Corallococcus* species revealed a predominance of GT enzymes, followed by GH, CBM, CE, AA, and PL enzymes. These enzymes were further categorized into 20, 40, 24, 9, 8, and 9 subcategories, respectively. Notably, eleven CAZymes, including CBM4, CBM56, CBM63, CBM9, CE11, GH128, GH44, GT19, GT30, GT35, and GT5, were found as single-copy genes in all examined species. GT4 emerged as the most abundant CAZyme overall **(Fig. 5)**.

**Fig. 5.**
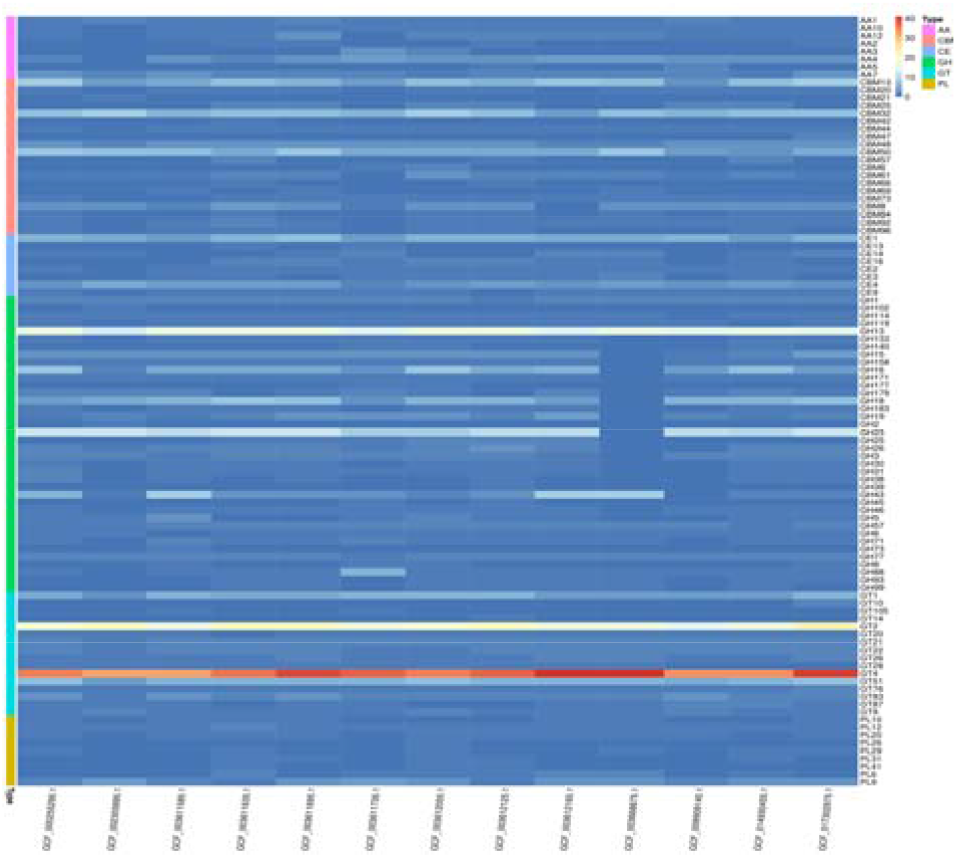
Heatmap representation of CAZY enzymes abundance profiles in *Corallococcus* species genomes.

## 4. Discussions

For many years, biotechnology and medicine have found great resources in nature’s chemical arsenal with an abundance of bioactive natural chemicals and their secondary metabolites. These diverse and potent molecules have served as the foundation for countless therapeutic agents for treating and preventing a wide range of human diseases [34]. However, with the expansion of genomic data, the domains of synthetic biology and genome mining are coming together to form a potent alliance that will unlock mysterious biosynthetic pathways and turn them into new drugs. By using NCBI’s publicly available genomic data for *Corallococcus* species and *Corallococcus exiguuus* subspecies reference strains, this study tried to find a lot of possible drug-like metabolites. Our results demonstrated that *Corallococcus* species and *C. exiguuus* subspecies harbor a vast natural product metabolic potential, characterized by high diversity. To ensure a reliable and well-characterized foundation for our study, we opted to exclusively utilize NCBI Reference Sequence (RefSeq) data. In our study, all 13 studied species exhibited distinct genomic signatures, based on the analysis of ANI and dDDH values. The observed GC content of 70.15% at the species level and 69.59% at the subspecies level suggests that, these organisms possess a genetic foundation for secondary metabolite production [24]. We observed a range of 2.56% to 4.30% of genes involved in secondary metabolite (SM) biosynthesis, transport, and catabolism across different *Corallococcus* species which indicates secondary metabolite production is a common and potentially crucial feature for these species. The absence of acquired antibiotic resistance genes and pathogenicity renders *Corallococcus* species attractive for potential and safe applications in biotechnology. This study aimed to elucidate the genetic basis of bioactive and CAZY enzyme production in *Corallococcus* species and *C. exiguus* subspecies. We observed a correlation between genome size and the number of biosynthetic gene clusters (BGCs) in *Corallococcus* species, suggesting a potential link between genomic complexity and enzyme diversity while no such association was found in *C. exiguus* subspecies. The distribution of biosynthetic gene clusters (BGCs) differed between the species and subspecies levels. At the species level, there were a total of 613 BGCs, encompassing 22 distinct types. In contrast, the subspecies level harbored 601 BGCs, representing only 16 types. This observation suggests a greater diversity of BGCs at the species level compared to the subspecies level. NRPS was predominant at both the species and subspecies levels, constituting 31.6% and 25.6% of the identified BGCs, respectively. The presence of a substantial number of completely unknown BGCs, observed at both the species and subspecies levels, hints at their potential as a reservoir for novel antimicrobials and bioactive products with promising applications in various sectors.

In our study, we found both *Corallococcus* species and *C. exiguus* subspecies to be valuable sources of antimicrobial peptides. Notably, both harbor lanthipeptide class II biosynthetic gene clusters, encoding core peptides with antimicrobial activity. This finding opens the doors for further research into the specific lantibiotics produced by these *Corallococcus* strains and their potential applications. On the basis of the presence of antibiotic-resistant target genes, twenty-one NRPS BGCs were prioritized as follows: *C. carmarthensis* (2), *C. exercitus* (2), *C. exiguus* (1), *C. interemptor* (1), *C. llansteffanensis* (5), *C. praedator* (2), *C. sicarius* (3), *C. soli* (1), and *C. terminator* (4). It is worth noting that it was predicted that these prioritized BGC regions would have the capacity to generate novel secondary metabolites that exhibit promising antimicrobial properties. Additionally, our analysis revealed that, among all the *Corallococcus* species that were examined, *C. llansteffanensis* demonstrates the most anticipated potency in generating novel secondary metabolites possessing antimicrobial properties. Investigation of the CAZymes repertoire in *Corallococcus* strains demonstrated the dominance of GT enzymes, followed by the presence of GHs. Notably, the representation of CBMs, CEs, AAs, and PLs further indicates their diverse metabolic capabilities. This enzymatic profile suggests the potential of *Corallococcus* strains to contribute significantly to human health.

## 5. Conclusions

In summary, we looked into the complete genomic makeup, secondary metabolite biosynthesis potential, polymer-degrading enzyme capabilities, and unique genomic features of *Corallococcus* at species and subspecies level. The strains of this marine bacteria harbor a diversity of biosynthetic gene clusters (BGCs), suggesting the potential to produce novel and valuable compounds. Further exploration of these prospective BGCs is needed to unveil their functional roles and potential applications. Additionally, *Corallococcus* possesses the ability to degrade a wide range of polymers, and has demonstrated the potential to fulfill the diverse nutritional requirements of crop plants, enhancing their growth and productivity. Moreover, *Corallococcus* exhibits the ability to produce compounds with potential therapeutic properties, paving the way for its utilization in the development of novel pharmaceuticals and nutraceuticals.

