## Supplementary figures and images for "Exploration of the Metabolic Potential of the *Corallococcus* Genus: A Rich Source of Secondary Metabolites, and CAZymes"

### aWlGINIQZmXVtGFuZyY45Q.png

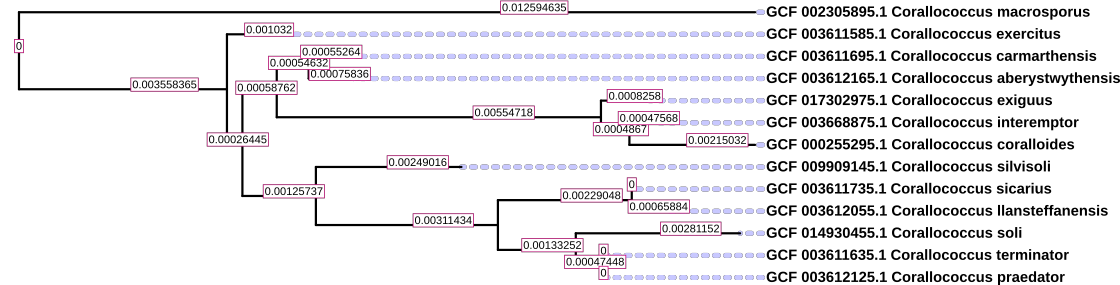

### ge.png

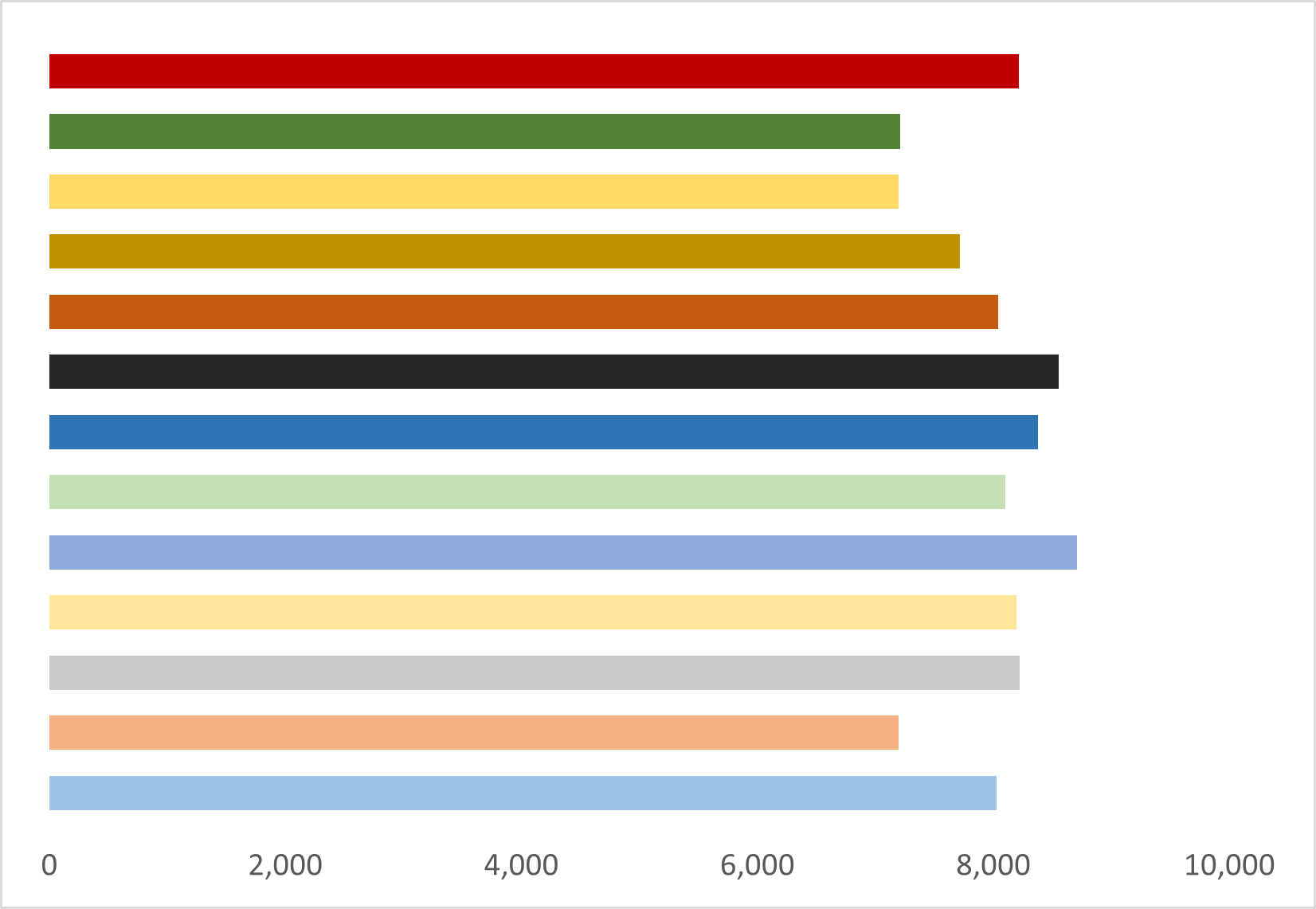

### phy_each_draft.png

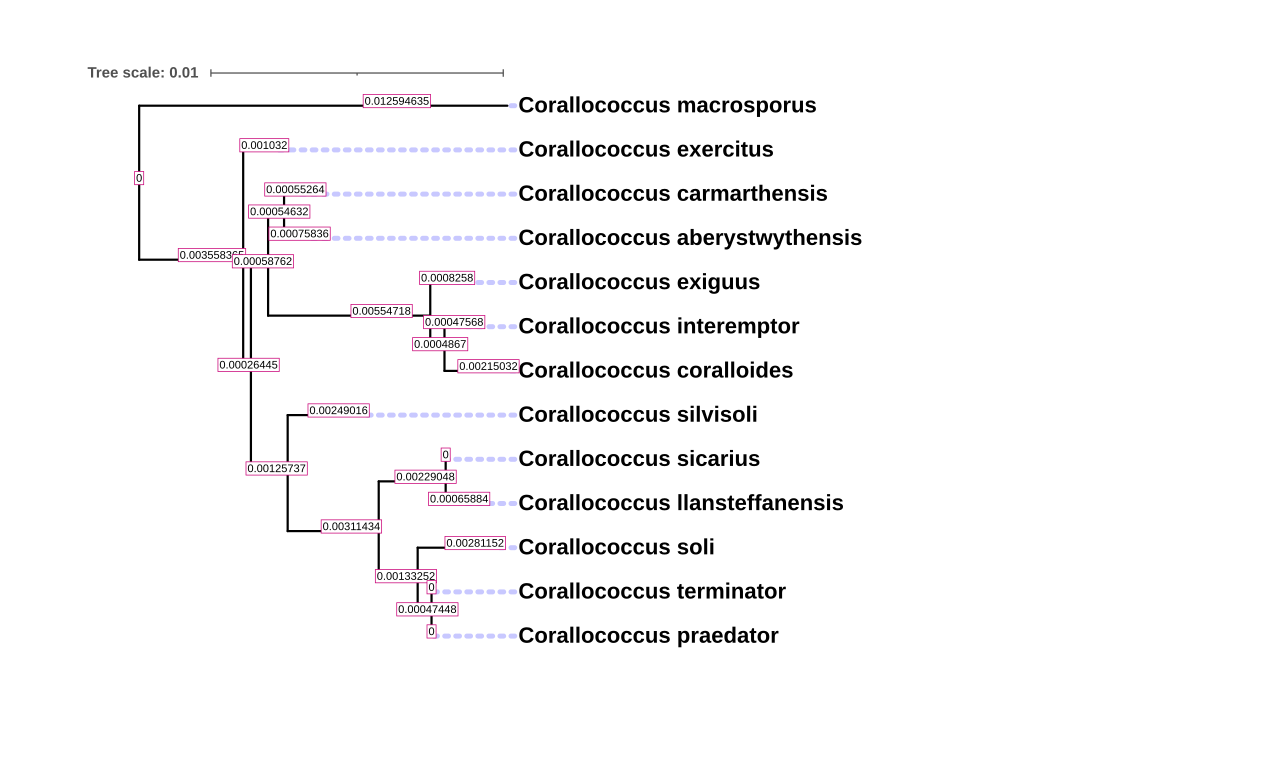

### phy_each_draft_italic.png

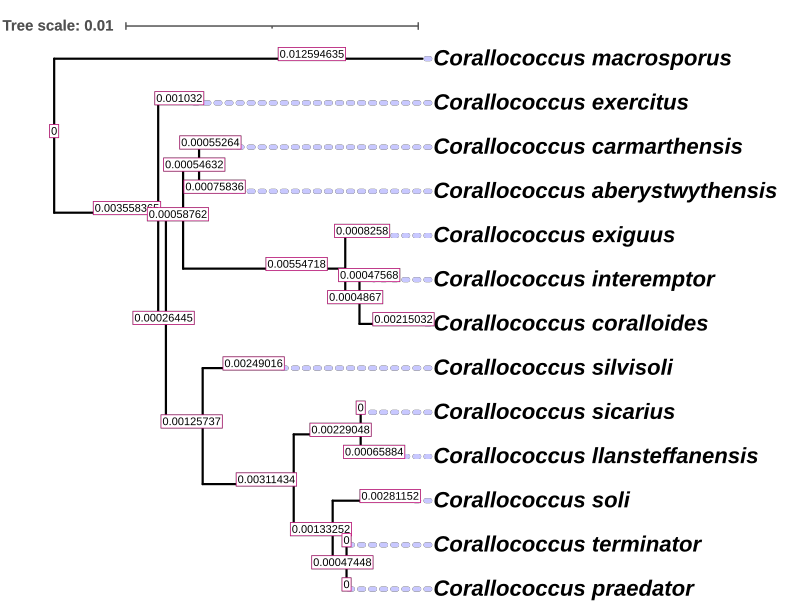

### phylo_draft_2.png

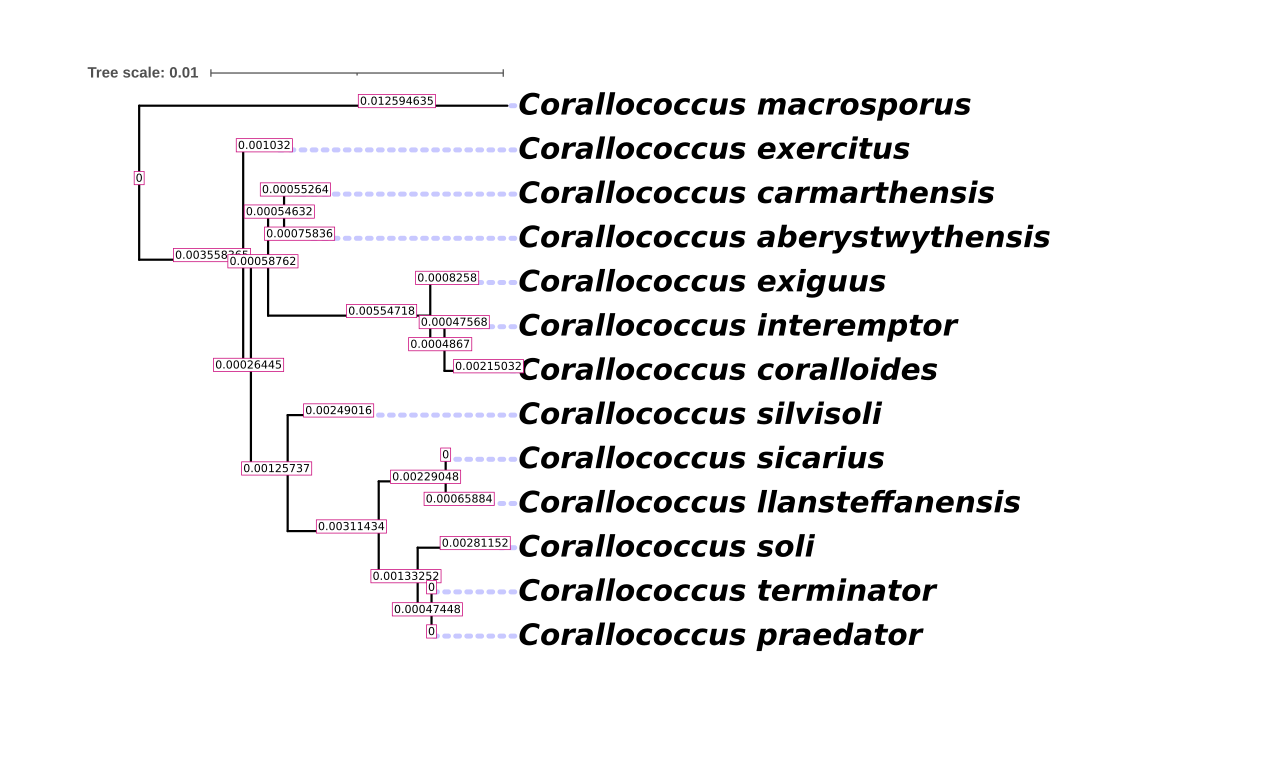

### Picture1.png

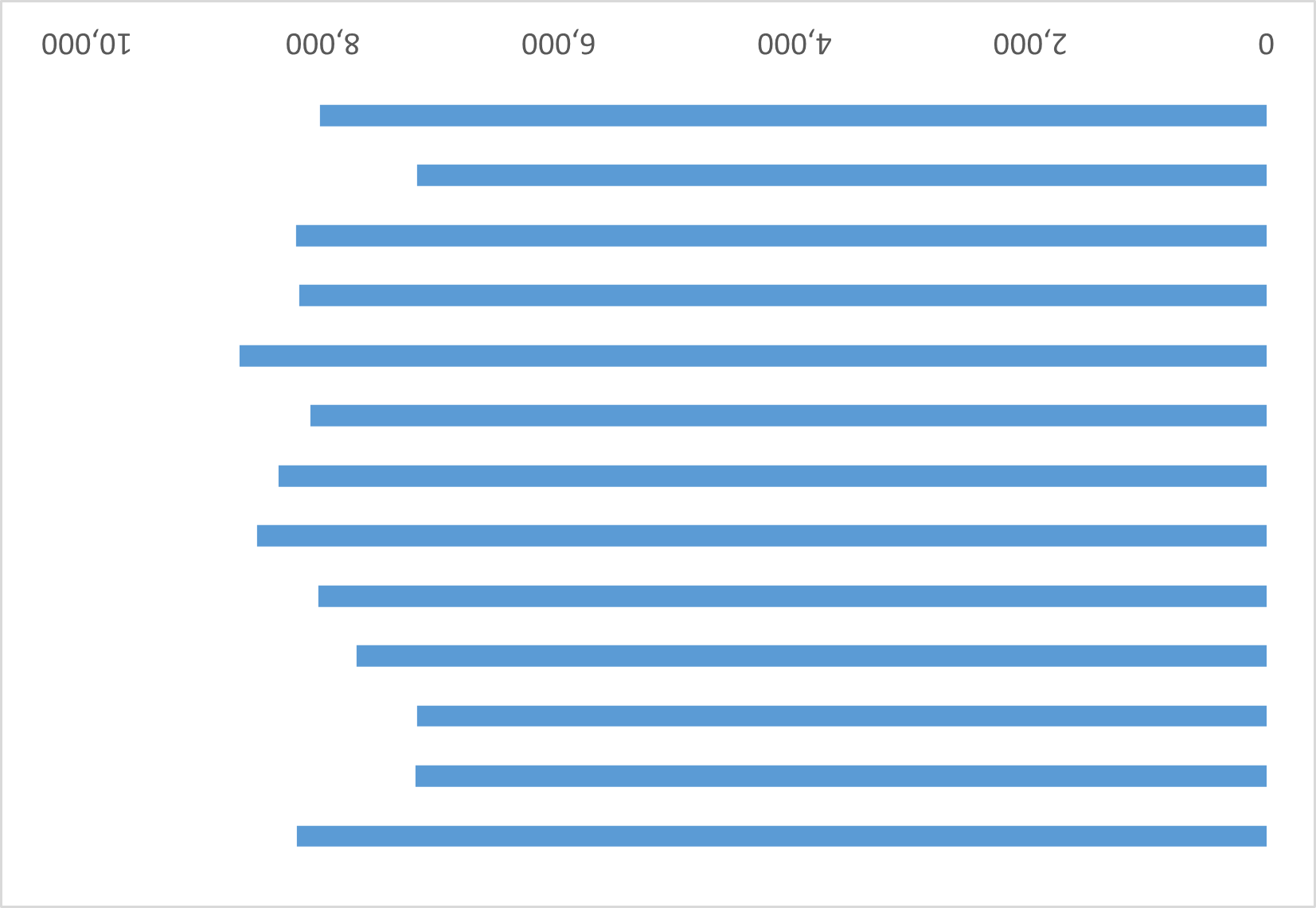

### Picture4.png

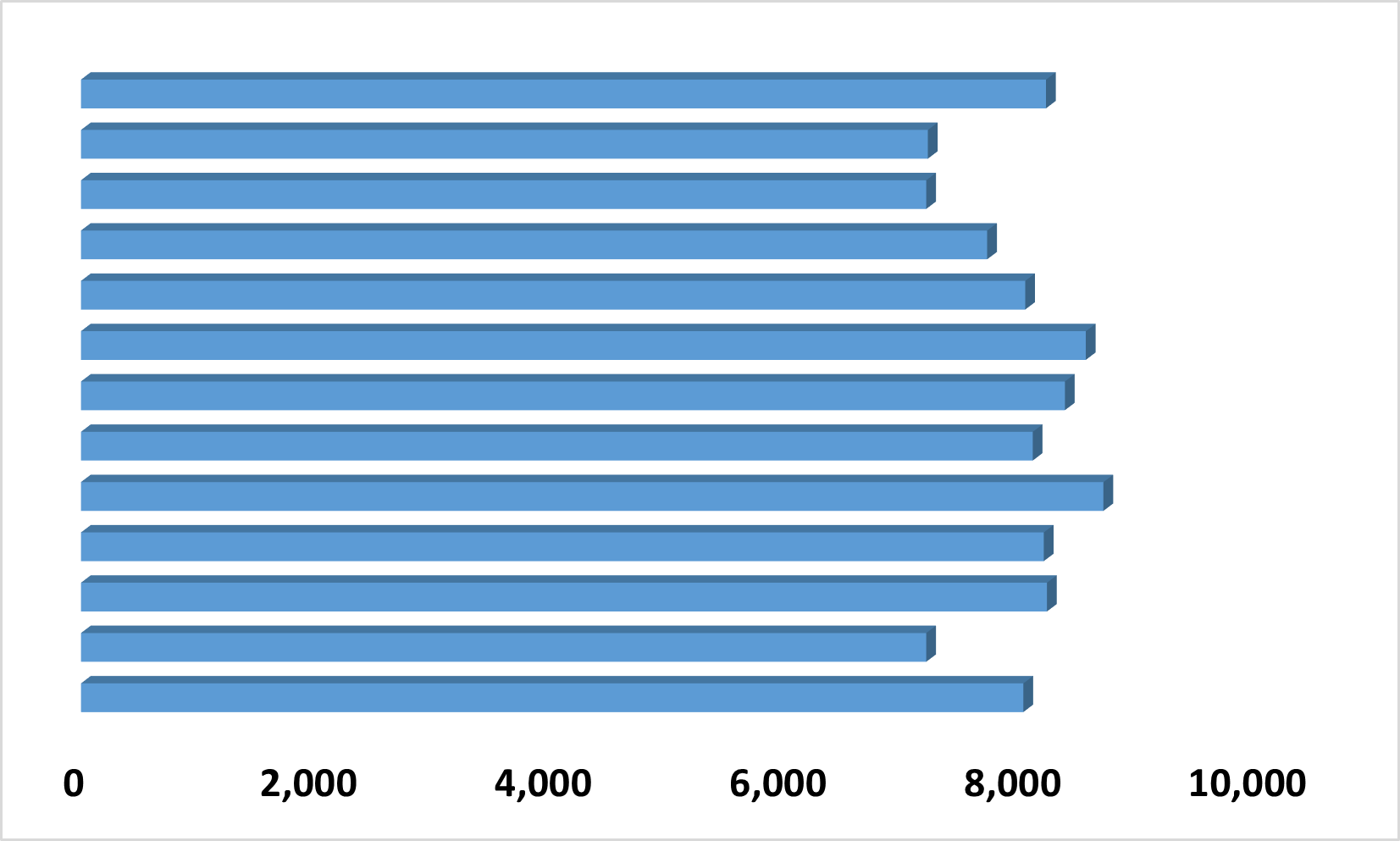

### Picture5.png

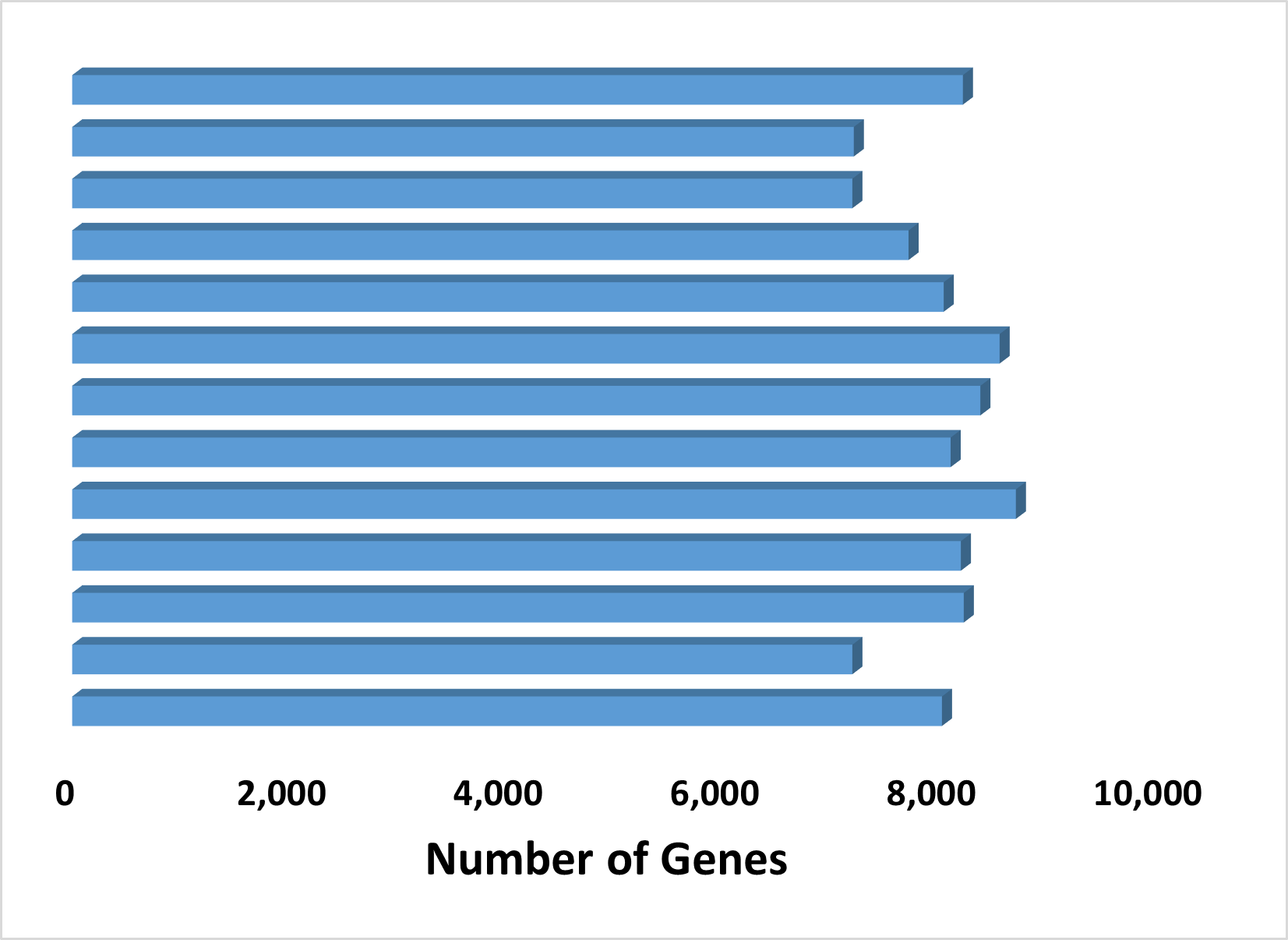

### Picture6.png

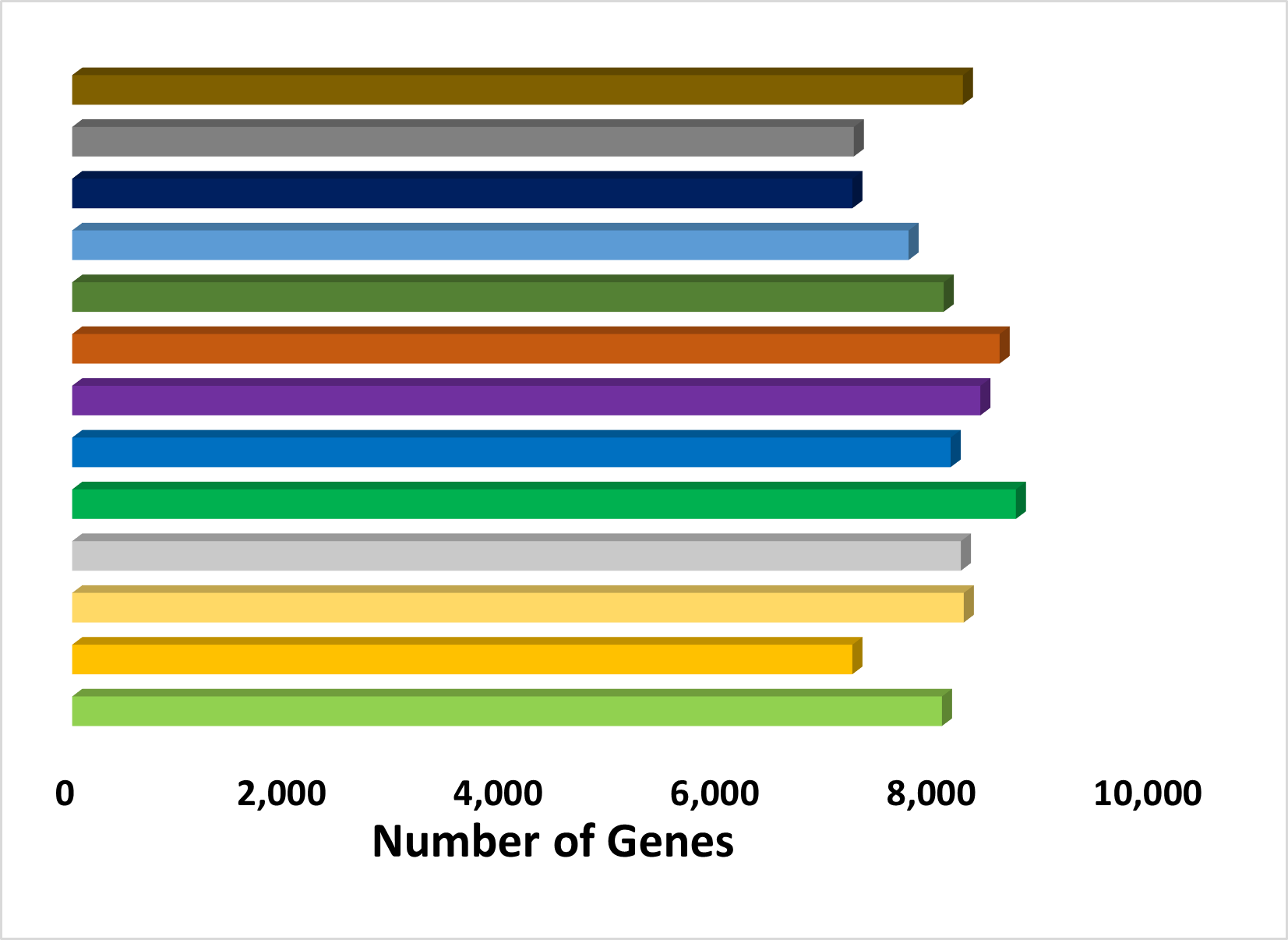

### Picture7.png

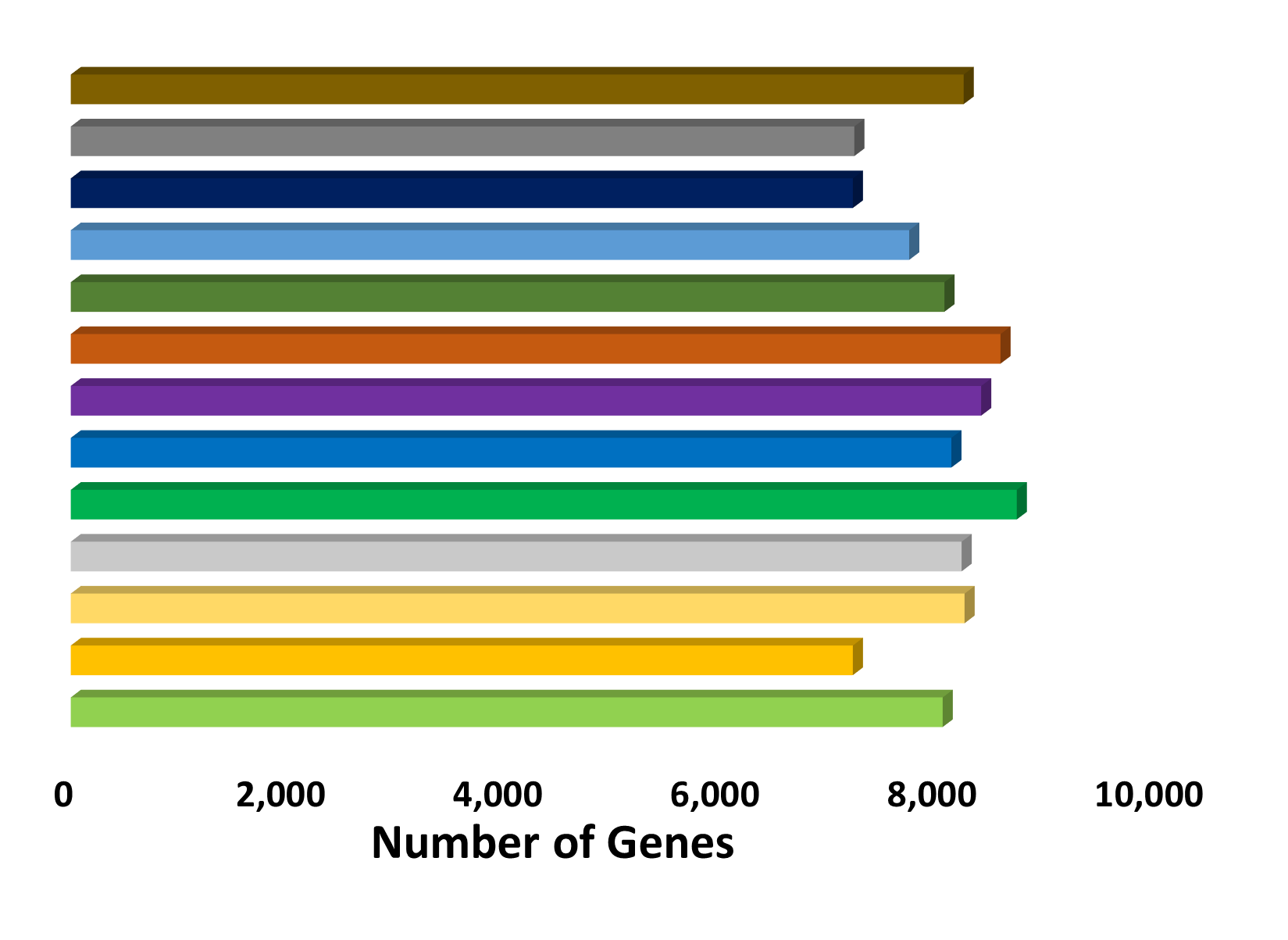
